# Eco-Friendly Antifouling Solutions: Hazard Assessment of Synthetic Derivatives of Natural Compounds

**DOI:** 10.64898/2026.03.31.715569

**Authors:** Jéssica Leite, Érica Lima, Daniela Pereira, Honorina Cidade, Marta Correia-da-Silva, Raquel Ruivo, Miguel M. Santos

## Abstract

The accumulation of microorganisms and macroorganisms on aquatic surfaces poses economic and ecological challenges, particularly in maritime transport. Traditional antifouling methods, such as biocidal coatings containing toxic compounds like tributyltin (TBT) and copper, are effective but harmful to the environment. This study investigates eco-friendly antifouling alternatives, focusing on nature-inspired compounds (NIAFs) GBA 26 (GBA) and DPC345DHC (DH345), derived from polyphenols and flavonoids, respectively.

The ecotoxicity of these compounds was evaluated using standardized assays with various species, including embryos of *Danio rerio* (zebrafish) (OECD TG 236), the algae *Raphidocelis subcapitata* (OECD TG 201), and the bacteria *Vibrio fischeri* (ISO 11348-2), along with nuclear receptor transactivation assays in *Mytilus galloprovincialis* (Mediterranean mussel). Gallic acid derivative GBA and 24h-transformation products showed low toxicity in zebrafish embryos, while dihydrochalcone DH345 inflicted developmental toxicity in zebrafish at 1 mg/L and above. Comparatively, tralopyril, a commercial biocide, exhibited significant toxicity at lower concentrations. Transcriptomic analysis of zebrafish embryos treated with GBA revealed selective gene modulation related to stress response, ion transport, and protein synthesis. Both, GBA and DH345, were shown to inhibit algae growth at 0.1 mg/L. *Vibrio fischeri* assay showed no toxic effects for any of the tested compounds. Nuclear receptor transactivation assays conducted with GBA revealed no activation of PPAR or PXR receptors.

These findings suggest GBA and DH345 as potential eco-friendly antifouling agents with lower environmental risks than established antifoulants such as tralopyril. However, further research is needed to evaluate their potential long-term ecological impacts, particularly chronic toxicity across various organisms. This study advances the pursuit of sustainable antifouling solutions that prioritize environmental protection.

## 1. INTRODUCTION

Microorganisms such as bacteria, protozoa, algae, and fungi readily colonize nutrient-rich solid surfaces, where they attach, proliferate, and form biofilms [1]. Over time, complex microbial consortia emerge, facilitating the settlement of larger organisms (macrofoulers), including macroalgae, mollusks, barnacles, and sponges [2, 3].

Biofouling is the unwanted accumulation of biofilms and associated organisms on surfaces in contact with liquid media.

It affects numerous industries, causing both economic and environmental burdens, particularly in maritime transport, by increasing hydrodynamic drag, fuel consumption by up to 40%, carbon dioxide emissions, and maintenance costs [4, 5].

Beyond economic concerns, biofouling promotes the spread of invasive species, disrupting local ecosystems and intensifying its environmental impact.

Antifouling coatings have long been used to reduce hull fouling and associated economic and environmental costs [6]. Among these, the organotin compound tributyltin (TBT) was widely applied for its high efficacy but was later banned due to its severe ecotoxicity. TBT caused endocrine disruption and imposex in gastropods such as *Nucella lapillus* [7, 8], along with growth inhibition, reproductive failure, and developmental abnormalities in fish and mammals [8–10].

Despite restrictions since the 90s [11], TBT persists in the environment due to its long half-life in water [12], entering food webs through bioaccumulation across trophic levels, ultimately posing risks to human health via contaminated seafood [13, 14].

Following its prohibition, alternative antifouling approaches were implemented using copper-based paints (e.g., CuSCN, Cu₂O), booster biocides (Irgarol 1051, Diuron, Chlorothalonil, Sea-Nine 211), along with copper-free biocides such as Econea (tralopyril), marketed as biodegradable [15]. However, such approaches remain problematic due to their specificity, which is often too narrow or too broad [16]. Additionally, the continuous release of copper and other biocides from antifouling paints raises concerns regarding their toxicity toward non-target species, leading to stricter environmental regulations and a push by the European Commission for safer, copper-free alternatives [17–19]. Hence, in the frame of previous projects, nature-inspired antifoulants (NIAFs) were synthesized and immobilized in commercial marine coatings [20]. While replacing copper-based systems is essential for sustainability, ensuring the antifouling efficacy and safety of these alternatives remains critical to avoid increased maintenance costs and ecological risks.

Recent studies have shown that certain NIAFs can effectively inhibit the settlement of *Mytilus galloprovincialis* larvae without evident toxicity to larvae or non-target organisms such as *Artemia salina* (<10% lethality at 25 µM), unlike the copper-free biocide tralopyril, which caused 100% lethality at the same concentration [21].

Among synthesized candidates, GBA, derived from molecular modification of a natural and affordable polyphenol, showed optimal potency against mussel larval settlement, with a superior LC50/EC50 ratio compared to tralopyril, maintaining efficacy when incorporated into polyurethane (PU)-based coatings [22]. GBA also reduced *Pseudoalteromonas tunicata* biofilms and decreased pre-existing ones [23].

Another compound, DPC345DHC (DH345), a synthetic dihydrochalcone, was obtained from the natural building block 2,4-dihydroxyacetophenone with 3,4,5-trimerhoxybenzaldehyde. DH345 showed to be highly active against the settlement of *M. galloprovincialis* larvae (EC_50_: 2.34 µM) with low toxicity against non-target marine species *A. salina* (< 1% at 25 and 50 µM). Moreover, when incorporated in a PU-based coating, DH345 maintained the anti-settlement activity, particularly at concentrations of 2% and 5% [24].

**Figure 1.**
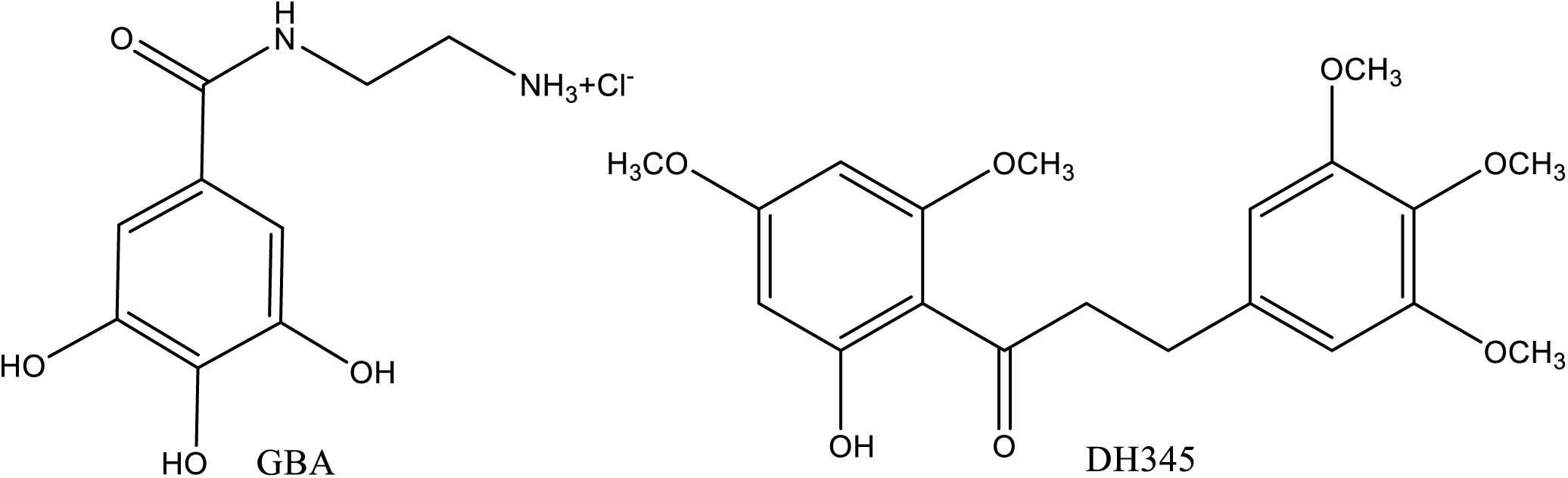
Synthetic antifouling compounds studied in this work. GBA= 2-(3,4,5-Trihydroxybenzamido)ethan-1-aminium chloride; DH345= 1-(2-hydroxy-4,6-dimethoxyphenyl)-3-(3,4,5-trimethoxyphenyl)propan-1-one

In the European Union (EU), the introduction of new biocides is regulated by the Biocidal Products Regulation (BPR, Regulation (EU) 528/2012) and REACH (Regulation (EC) No. 1907/2006), both of which are overseen by the European Chemicals Agency (ECHA). These frameworks require comprehensive risk assessments, including hazard assessment, exposure assessment, and risk characterization, to ensure the safe use of chemicals across the supply chain. For antifouling products, an Environmental Risk Assessment (ERA) must be performed prior to market approval. This involves estimating Predicted Environmental Concentrations (PECs) and Predicted No-Effect Concentrations (PNECs) to derive risk characterization ratios. Supporting data, such as LOEC, NOEC, and ECx values, are essential for establishing PNECs and defining regulatory safety limits [25].

Manufacturers must submit a detailed dossier to ECHA, including data obtained under the Organization for Economic Cooperation and Development (OECD) Test Guidelines, demonstrating that the product is both effective and safe for human health and the environment. This integrated regulatory approach ensures that only biocides meeting high safety and environmental standards are authorized for use within the EU [26].

This study aims to assess the ecotoxicological impact of these novel antifouling NIAFs across different species using standardized tests. Given that one major challenge in handling chemical compounds lies in their potential to act as endocrine-disrupting chemicals (EDCs), their potential endocrine-disrupting activities were also evaluated using Nuclear Receptors as a proxy. Nuclear receptors (NRs), a superfamily of ligand-activated transcription factors, play a central role in mediating endocrine and environmental signals by regulating gene expression involved in metabolism, reproduction, immunity, and development [27].

Due to their ligand-activated nature, NRs are prime targets for EDCs, and may lead to diverse health and environmental consequences. Consequently, NRs serve as valuable tools for screening potential EDCs [5], and play a key role in the OECD guidelines for the evaluation of endocrine disruptors [28–32].

By examining the effects of GBA and DH345 across various trophic levels, this study aims to improve hazard and risk assessments, contributing to the development of sustainable antifouling solutions.

## 2. MATERIAL AND METHODS

### 2.1. Chemicals

Hydrochloride of GBA 26 (GBA) and DPC345DHC (DH345) were provided by the Faculty of Pharmacy of the University Porto [24, 33]. ECONEA® (CAS no: 122454-29-9, Janssen PMP), dimethyl sulfoxide (DMSO; CAS no: 67-68-5, SupraSolv®) was used as a solvent, RNAlater® (Sigma), dechlorinated water (CIIMAR), a Microtox® kit (AmbiFirst), and an RNA extraction kit (NZYtech).

### 2.2. Fish Embryo Acute Toxicity Test

This study followed OECD Test Guideline No. 236, “Fish Embryo Acute Toxicity (FET)” [34]. Zebrafish (*Danio rerio*) embryos, AB strain, were exposed to GBA (10, 1, 0.1 mg/L), DH345 (1, 0.7, 0.35, 0.1, 0.01 mg/L), and tralopyril, as a positive control (5, 2.27, 1.03 µg/L), plus water control.

Twenty fertilized embryos (≈1.5 hours post fertilization (hpf)) per treatment were individually placed in 24-well plates and maintained for 144 hpf at 26 ± 0.5 °C under a 14:10 light-dark cycle, with daily medium renewal. Mortality and developmental abnormalities, such as oedema, growth arrest, and morphological defects, were recorded daily. Larval morphometrics (yolk sac, eye area, body length) were assessed at 48, 96, and 144 hpf using a digital camera and an inverted microscope (Nikon Eclipse TS100). After exposure, larvae were stored in RNAlater at –80 °C for RNA extraction. The assay was replicated three times

#### 2.2.1. Degradation of GBA in 24 hours-FET samples

Considering the high degradability in water, GBA concentration was quantified during FET assay.

A chromatographic method was defined to quantify GBA. HPLC system consisted of a pump, autosampler, column compartment, variable wavelength detector and fluorescence detector, all from Dionex Ultimate 3000 (Thermo Fisher Scientific, Bremen, Germany). The software used to control the system and perform data treatment was Chromeleon 7. The method consisted of mobile phase (MP) A of water acidified with 0.1% formic acid and MP B of acetonitrile. Run time of 10 min in isocratic mode: 50:50, v:v, MP A: MP B. The stationary phase was the column Kinetex 1.7 μm EVO C18 100 Å, 100 x 2.1 mm (Phenomenex, California, USA) at room temperature. Chromatographic conditions were set at flow rate of 0.1 mL/min, injection volume of 1.0 μL and UV detection wavelength of 267 nm. A calibration curve was performed with six concentrations of GBA (5, 10, 20, 30, 40 and 50 μg/mL) in dechlorinated water (Supplementary Material).

Aiming to identify the transformation products (TPs) of GBA, a sample (dechlorinated water) exposed to the ecotoxicity assay for 24 hours was analyzed using liquid chromatography coupled with high-resolution mass spectrometry (LC-HRMS). Additionally, freshly prepared standards were analyzed in dechlorinated water. The analyses were conducted at the Materials Center of the University of Porto (CEMUP) using an Orbitrap Exploris 120 mass spectrometer (Thermo Fisher Scientific, Bremen, Germany), operated via the Orbitrap Exploris Tune Application version 2.0.185.35 and Xcalibur software version 4.4.16.14. The electrospray ionization (ESI) source was configured with a capillary voltage of 3.4 kV for positive mode and 2 kV for negative mode, while the capillary temperature was maintained at 320°C. Sheath and auxiliary gas flow rates were set to 5 arbitrary units, as defined by the software. The resolution for MS scans was 60,000. Data-dependent MS/MS was carried out in HCD mode using nitrogen as the collision gas, with a collision energy of 30 V. The m/z range was set between 100 and 1000 Da. The resolution for SIM MS scans was also 60,000. The identification of TPs was performed by matching their m/z values and MS/MS spectra using the Xcalibur QualBrowser software (Thermo Fisher Scientific). The method utilized a mobile phase (MP) composed of water with 0.1% formic acid (MP A) and acetonitrile (MP B). The analysis was performed in isocratic mode with a 10-minute runtime, using a 50:50 (v/v) ratio of MP A to MP B. The stationary phase was a Kinetex EVO C18 column (1.7 μm particle size, 100 Å pore size, 100 × 2.1 mm; Phenomenex, California, USA), maintained at room temperature. Chromatographic parameters included a flow rate of 0.1 mL/min, an injection volume of 10 μL, and UV detection at a wavelength of 267 nm.

#### 2.2.2. Transcriptome Analysis of zebrafish embryos

Based on the outcome of zebrafish embryo bioassay, RNA-seq analysis was performed with the embryos from GBA, 0.1 mg/L treatment. Total RNA was extracted from zebrafish embryos exposed to GBA or water (control), with four replicates per condition (pool of four embryos per replicate), and analyzed by Novogene. After RNA integrity validation, mRNA was isolated, converted to cDNA, and prepared for sequencing on the Illumina platform.

Raw reads were quality-filtered with *fastp* and mapped to the zebrafish reference genome using Hisat2 v2.0.5. Transcripts were assembled with StringTie v1.3.3b [35], and gene expression quantified as FPKM (Fragments Per Kilobase of transcript per Million mapped reads) using featureCounts v1.5.0-p3. Differential expression analysis was performed with DESeq2, identifying significantly differentially expressed genes (adjusted p ≤ 0.05; log2 fold change ≥ |1|).

Results were visualized in hierarchical clustering heatmaps based on normalized log2(FPKM+1) values and Z-scores were calculated. Functional enrichment of differentially expressed genes was assessed via clusterProfiler, identifying significantly enriched GO terms (p < 0.05) across Biological Process, Cellular Component, and Molecular Function categories.

### 2.3 Freshwater Alga Growth Inhibition Test

The freshwater alga growth inhibition test followed OECD Guideline No. 201 [36] using *Raphidocelis subcapitata* cultures (SKULBERG 1959; CCAP 278/4) maintained under controlled conditions: medium 3N-BBM+V [110] with constant agitation (200 rpm) and light (60-120 µmol/m²/s) under 21 ± 1 °C. Algae (10⁴ cells/ml/well) were exposed to GBA (10, 1, 0.1 mg/L), DH345 (0.35, 0.1, 0.001 mg/L) and DMSO (control) for 72 h in 24-well plates, and growth inhibition (%) was determined by microscope (Nikon Eclipse TS100) using a Neubauer chamber, relative to controls. The assay was replicated four times.

### 2.4 Vibrio fischeri Bioluminescence Assay

The Vibrio fischeri bioluminescence assay followed ISO 11348-2:2007 guidelines using the Microtox 500 Analyzer (Modernwater, New Castle, DE) according to the 81.9% Basic Test protocol [37]. Bacteria were exposed for 5, 10, and 30 min to serial dilutions of GBA (10 mg/L), DH345 (1 mg/L), and tralopyril (5 µg/L). Negative controls consisted of dechlorinated water and DMSO 0.01%; positive control consisted of 3,5-dichlorophenol 5 mg/L. Luminescence inhibition (EC20 and EC50) was calculated with MicrotoxOmni Azur® software.

### 2.5 Nuclear Receptor Transactivation Assays

These assays evaluated GBA capacity to activate nuclear receptor-mediated transcription of PPAR and PXR from *Mytilus galloprovincialis*. Fusion-proteins were obtained by combining hinge and ligand-binding domain (LBD) regions of the selected nuclear receptors with a yeast Gal4 DNA-binding domain (DBD) which acts on activation sequences (UAS) upstream a firefly luciferase reporter gene [38]. *M. galloprovincialis* receptor PPAR [Accession: MGAL_10B094094] was kindly provided by Angelica Miglioli (Centre National de la Recherche Scientifique (CNRS), Villefranche-sur-Mer, France) [39]; PXR [40] was kindly provided by Elza Fonseca (Interdisciplinary Centre of Marine and Environmental Research (CIIMAR)). COS-1 cells (ATCC, USA) were cultured at 37°C, 5% CO₂ in DMEM with 10% FBS and 1% penicillin/streptomycin (PAN-Biotech, Aidenbach, Germany). Cells (2×10⁵/mL/well) were seeded in 24-well plates and, after 24 h, transfected with 0.5 μg of reporter vector (pGAL4.35) and 0.5 μg of receptor constructs (PPAR/pBIND or PXR/pBIND) using Opti-MEM (Gibco, USA), Lipofectamine® 2000 (Invitrogen, USA), and incubated for 5 h. After rinsing with Phosphate

Buffer Saline (PBS; PAN-Biotech, Aidenbach, Germany), cells were exposed for 24 h to GBA (0.1, 1, and 10 µM). Firefly and Renilla luciferase activities were quantified using the Dual-Luciferase Reporter Assay System (Promega, USA) and a Synergy HT Multi-Mode Microplate Reader (BioTek, USA).

### 2.6 Statistical Analyses

Data was analyzed using IBM SPSS Statistics 29.0.0.0. Variance homogeneity was tested (Levene’s test), followed by one-way ANOVA and Dunnett’s post-hoc test (α = 0.05). If variances were non-homogeneous, the Kruskal-Wallis test with Bonferroni correction was applied.

## 3. RESULTS AND DISCUSSION

### 3.1 Degradation of GBA during FET assays

Knowing the high degradability of GBA in seawater, GBA samples that were incubated 24h in the FET medium were analyzed by HPLC-UV and LC-MS. This property makes GBA particularly attractive as an antifouling biocide, as its quick breakdown reduces the potential for long-term environmental contamination, bioaccumulation, and trophic transfer.

After developing the calibration curve in dechlorinated water (Figure S1), GBA in dechlorinated water exposed for 24h in the FET assay was quantified (Figure 3).

**Figure 2.**
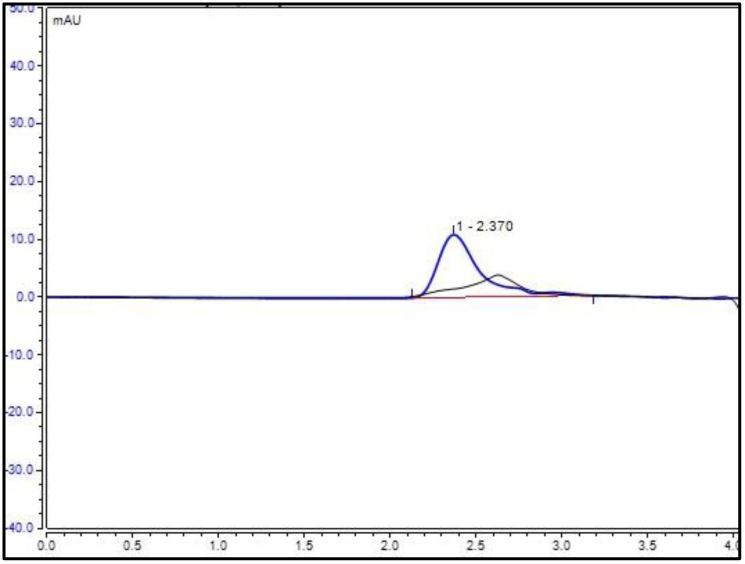
Representative chromatogram of GBA after 24h in dechlorinated water in FET assay and standard freshly prepared (blue,10 µg/mL).

Following, LC-HRMS was used to analyse the sample of GBA after 24h in dechlorinated water in FET assay. In the Supplementary material, the chromatogram and mass spectrum (positive ionization mode) of a freshly prepared GBA standard in dechlorinated water are presented (Figure S2). GBA appears at a retention factor between 1.64-2.20 seconds with a mass-to-charge ratio (m/z) of 213.086, representing the molecule without the chloride ion and with an additional proton from positive ionization. In Figure S3, the chromatogram and mass spectrum of the 24 hours-FET sample in dechlorinated water are shown. The mass-to-charge ratio (m/z) of 213.086 cannot be found (Figure 4) .

**Figure 3.**
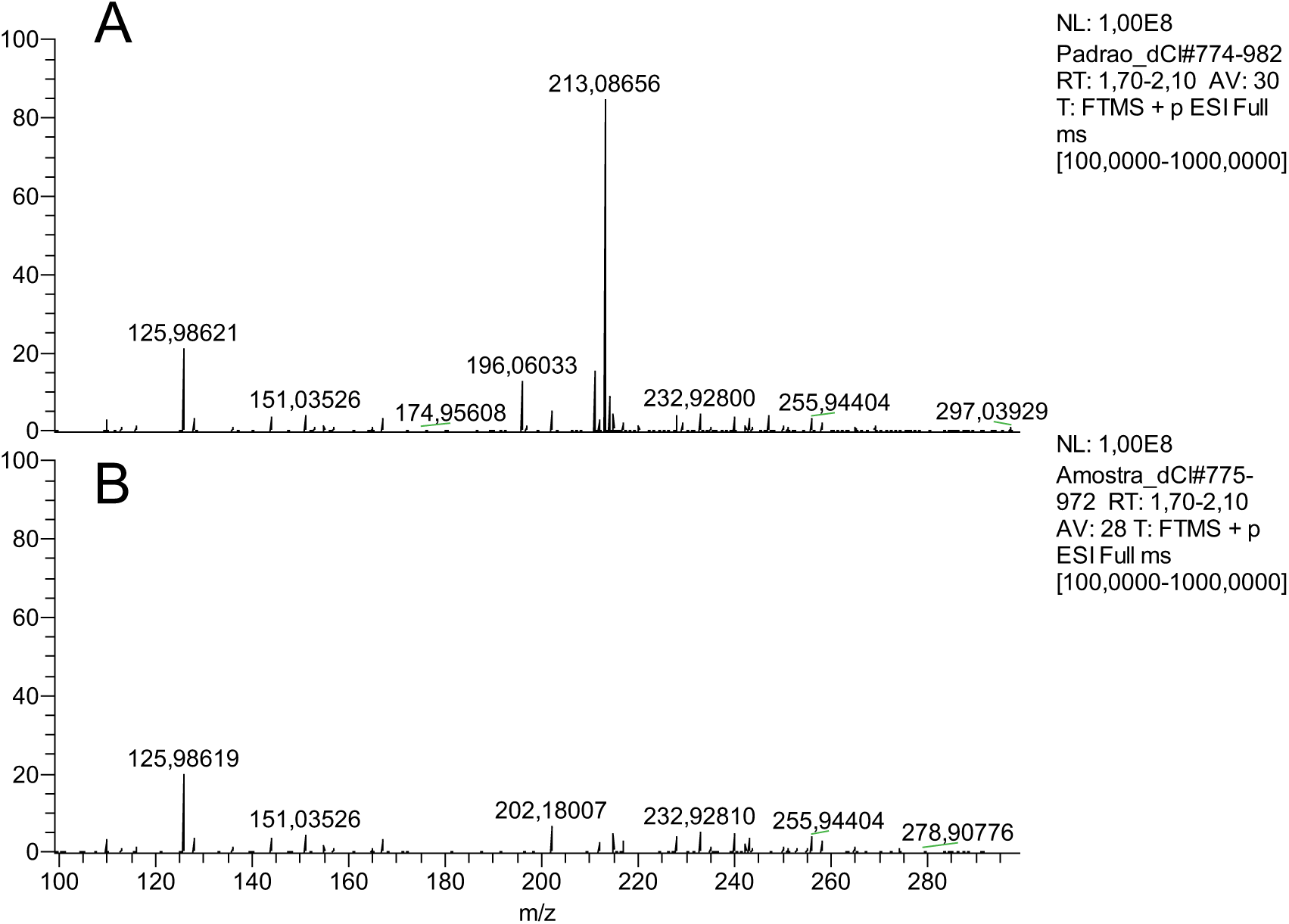
Mass spectrum of GBA freshly prepared (A) and GBA after 24h in dechlorinated water in FET assay (B).

These results underscore the importance of daily renewal of test solutions to maintain experimental consistency and ensure that the observed effects were due to GBA, given the compound’s rapid degradation rate.

### 3.1 Fish Embryo Acute Toxicity Test

Fish embryo toxicity (FET) tests were carried out to evaluate the potential hazard of GBA and DH345.

All assays met the OECD control criteria, with ≥90% survival and normal development. No significant differences were found between dechlorinated water and DMSO controls for any endpoint, so they were merged into a single control for data treatment. DMSO was used only for tralopyril and DH345 treatments, while GBA was tested using dechlorinated water due to its high water solubility.

**Figure 4.**
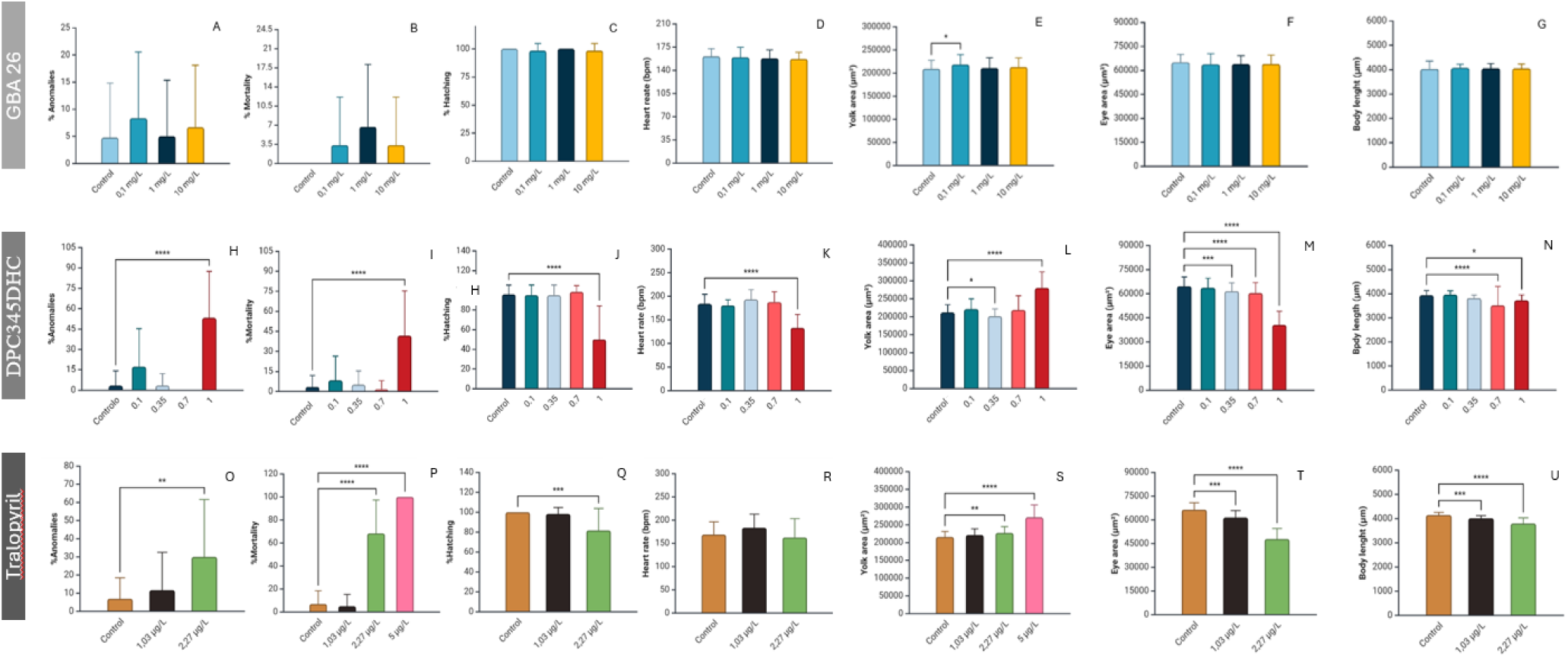
Effects of GBA exposure (Control, 0.1 mg/L, 1 mg/L, and 10 mg/L), DPC exposure (Control, 0.1 mg/L, 0.35 mg/L, 0.7 mg/L and 1 mg/L) and tralopyril exposure (Control, 1.03 μg/L, 2,27 μg /L, and 5 μg /L) on different developmental and physiological parameters. (A, H, O) Percentage of anomalies (measured at 144 hpf) (B, I, P) Percentage of mortality (measured at 144 hpf) (C, J, Q) Percentage of hatching (measured at 144 hpf) (D, E, R) heartbeat rate (bpm) (measured at 72 hpf) (E, L, S) Yolk sac area (µm²) (measured at 48 hpf) (F, H, T) Eye area (µm²) (measured at 96 hpf) (G, N, U) Body Length (µm) (measured at 144 hpf). (A-N) One-way-ANOVA (p&<0,05) with Dunnett post-hoc test. Error bars represent standard deviations. * Indicates statistically significant differences in relation to the control (p &< 0.05). (O-U) Krustal-Wallis Test (p&<0,05) with Bonferroni correction. Error bars represent standard deviations. * Indicates statistically significant differences in relation to the control (p &< 0.05); ** (p &< 0.01); *** (p &< 0.001); **** (p &< 0.0001).

Three different assays were conducted, one per compound (GBA, DH345, tralopyril). Each test assessed multiple endpoints: percentage of anomalies, mortality, hatching success, heart rate (bpm), yolk sac area (µm²), eye area (µm²), and body length (µm).

Figure 4A–G illustrates the results for GBA exposure. Overall, no statistically significant differences were found between treated and control groups for anomalies, mortality, hatching, heart rate, eye area, or body length. A statistically significant increase (p < 0.05) was only observed in yolk sac area, at 0.1 mg/L, which was not replicated at higher concentrations (1 and 10 mg/L), indicating the absence of a clear dose-response trend, possibly explained by a hormetic dose-response pattern [41].

Overall, exposure to GBA caused minimal alterations in most parameters, suggesting no significant developmental or toxic effects on zebrafish embryos. The stability of endpoints such as hatching, growth, and especially heart rate supports the conclusion that GBA did not induce physiological stress or disrupt normal embryonic development [42].

The effects of DH345 at 0.1, 0.35, 0.7, and 1 mg/L on zebrafish embryonic development and physiology are shown in Figure 2H-N.

At 1 mg/L, a significant increase in anomalies was observed (Figure 4H), while lower concentrations showed no differences from the control. Mortality (Figure 4I) also rose significantly at 1 mg/L but remained unchanged at lower doses. Similarly, hatching rate (Figure 4J) was significantly reduced at 1 mg/L, whereas no alterations were observed in the other treatments. Heart rate (Figure 4K) was significantly lower in embryos exposed to 1 mg/L, but unaffected at lower doses, indicating cardiac sensitivity only at the highest concentration.

In the yolk sac area (Figure 4L), a significant decrease was observed at 0.35 mg/L, whereas an increase was observed at 1 mg/L, suggesting opposite responses at these concentrations. Eye area (Figure 4M) progressively decreased in 0.35, 0.7, and 1 mg/L groups, with no changes at 0.1 mg/L. Body length (Figure 4N) was significantly reduced at 0.7 and 1 mg/L, but not at lower doses, indicating that growth inhibition occurs at higher exposures.

The pronounced increase in anomalies and mortality at 1 mg/L suggests that DH345 may interfere with key developmental processes (e.g. cell proliferation or differentiation). The corresponding decline in hatching success supports this hypothesis, implying that embryos exposed to high DH345 concentrations fail to complete the developmental programme.

Unregulated cell growth and differentiation may also explain the reduction in overall larval growth. Additionally, a lower heart rate at the highest dose further indicates that DH345 may impair cardiovascular function.

Altogether, these findings demonstrate a clear dose-dependent response: low concentrations produce negligible effects, whereas higher concentrations lead to significant developmental and physiological disruptions. Similar to GBA, the precise mode of action (MoA) of DH345 remains unknown, and further research is needed to clarify its biological targets and mechanisms. The effects of tralopyril at 1.03, 2.27, and 5 μg/L on zebrafish development and physiology are shown in Figure 4O-U.

At 5 μg/L, all larvae died by 72 hpf, allowing only mortality and yolk sac data collection. A significant increase in anomalies was observed at 2.27 μg/L compared to the control (Figure 4O). Mortality rose significantly at both 2.27 μg/L and 5 μg/L (Figure 4P), indicating severe toxicity at higher concentrations. Hatching rates decreased at 2.27 μg/L (Figure 4Q), while heart rate remained largely unaffected (Figure 4R). Yolk sac area increased significantly at 2.27 μg/L and 5 μg/L (Figure 4S), suggesting disrupted yolk absorption or developmental delay. Eye area decreased notably with increasing dose, especially at 2.27 μg/L (Figure 4T). Body length was also significantly reduced at 2.27 μg/L, with a milder reduction at 1.03 μg/L (Figure 4U). The absence of data for the 5 μg/L group in some parameters reflects the complete mortality at this concentration. These results demonstrate that tralopyril exerts severe toxic and developmental effects at higher doses.

These findings highlight the high toxicity potential of tralopyril to aquatic organisms, consistent with previous studies reporting its severe effects on fish development and survival [43].

### 3.2 Transcriptomic Analysis of GBA-exposed Zebrafish Embryos

Given the low toxicity of GBA and TPs observed in the apical endpoints, further analyses were performed to investigate its potential effects at a molecular level, particularly on gene expression and signalling pathways.

Figure 5A summarizes the transcriptional response of zebrafish embryos exposed to GBA and TPs, identifying 1860 differentially expressed genes (992 downregulated and 868 upregulated) relative to the H₂O control, indicating a clear molecular response despite the low apical toxicity observed. Hierarchical clustering of normalized gene expression (Figure 5B) revealed distinct and consistent expression patterns between treated and control groups, suggesting selective activation and repression of specific pathways rather than widespread transcriptomic disruption. Gene Ontology enrichment analysis (Figure 5C) showed a predominant downregulation of biological processes associated with protein synthesis and translation, including amide and peptide biosynthetic processes, consistent with a stress-related cellular response characterized by reduced global translation while maintaining the expression of selected protective or stress-related proteins [44].

**Figure 5.**
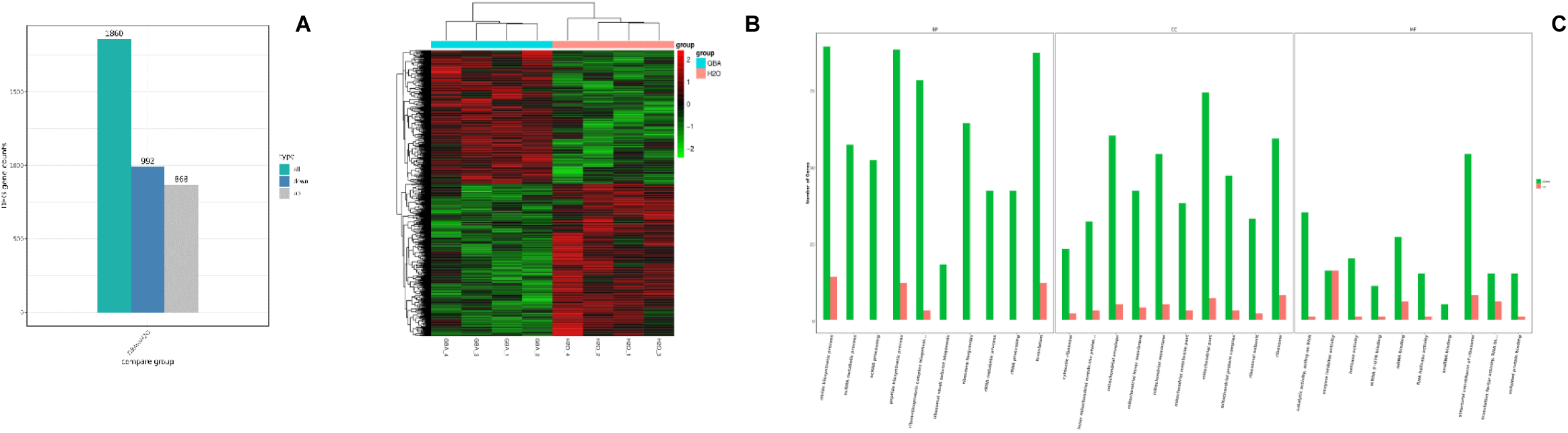
**(A)** Differentially Expressed Genes (DEG) between GBA and water control. The bar plot represents the total number of DEGs (all) as well as genes with significant downregulation (down) and upregulation (up) in response to GBA treatment compared to the water control. **(B)** Heatmap showing the hierarchical clustering of differentially expressed genes between zebrafish embryos treated with GBA 26 and the water control group. Each row represents a gene. Gene expression was analyzed by clustering the log2(FPKM+1) values. **(C)** GO Enrichment Analysis of Differentially Expressed Genes in GBA 26-Treated Zebrafish Embryos. This bar plot shows the number of upregulated (red) and downregulated (green) genes in enriched Biological Process (BP), Cellular Component (CC), and Molecular Functions.

Under stress conditions, cells typically inhibit global translation to prevent error-prone protein synthesis, while prioritizing the production of proteins essential for damage control and homeostasis [45]. The upregulation of these biosynthetic pathways could reflect the cell’s attempt to maintain the synthesis of critical proteins, even while downregulating overall protein production.

Principal component analysis (PCA; Figure 6) further supports these findings. While PC1 (37% of variance) does not clearly separate treated and control samples, PC2 (18% of variance) distinctly discriminates GBA and TPs–exposed embryos from controls, indicating specific transcriptional effects associated with the biocide rather than broad variability.

**Figure 6.**
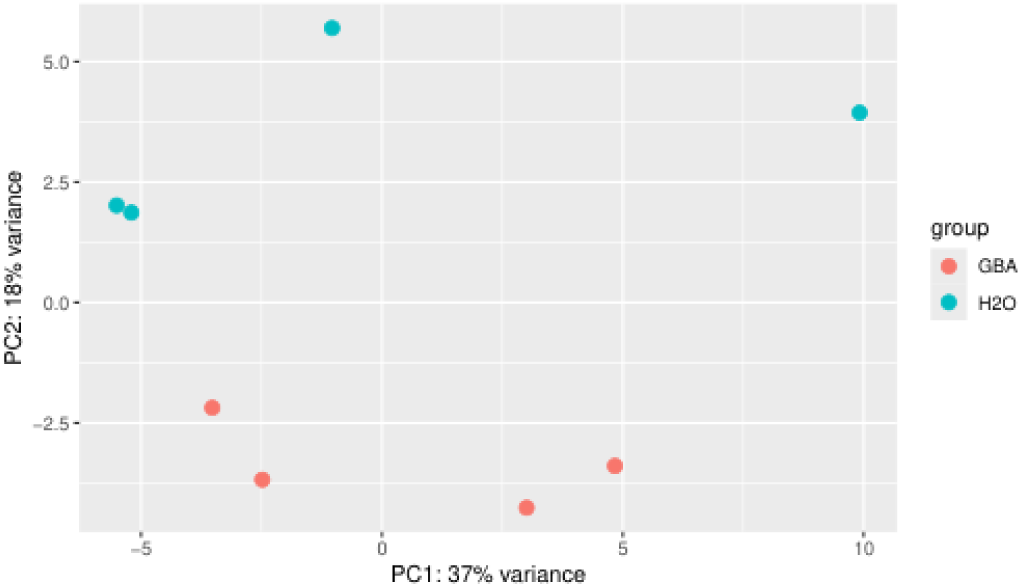
Principal component analysis (PCA) plot of gene expression data from zebrafish embryos treated with GBA 26 (red) and H2O control (blue). PC1 explains 37% of the variance and reflects general variability in the dataset, while PC2 explains 18%.

Several genes involved in stress response and cellular regulation were significantly altered (Figure 7; Table 1). Notably, *anxa1b* (downregulated) and *anxa1c* (upregulated), both encoding annexin proteins involved in inflammation, membrane repair [46], display opposite expression trends.

**Figure 7.**
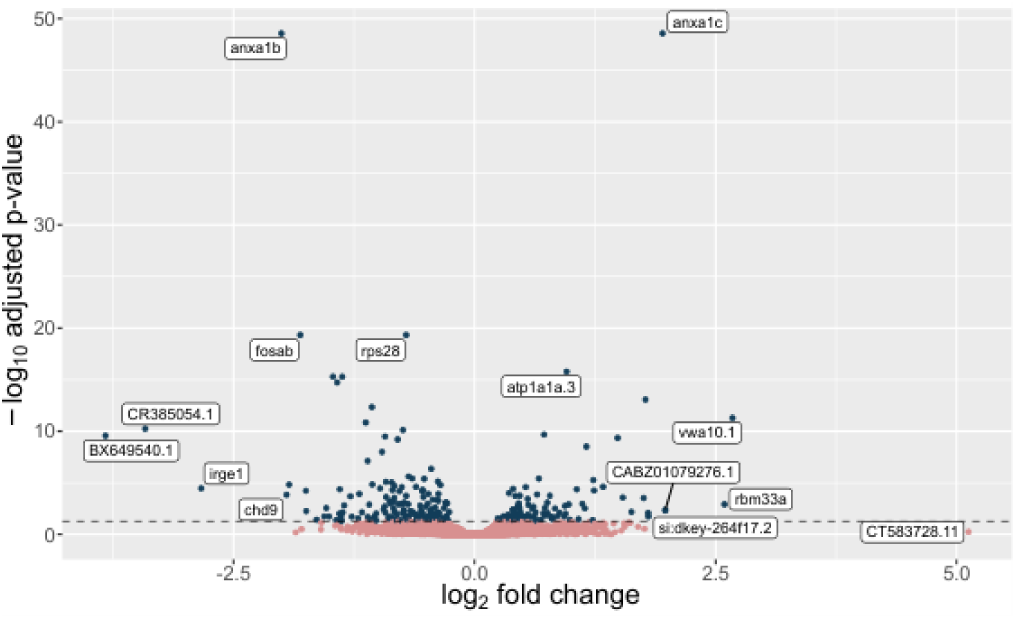
Volcano plot showing the differential gene expression between zebrafish embryos treated with GBA 26 and the H2O control group. The x-axis represents the log2 fold change, with positive/negative values indicating upregulated/downregulated genes in the GBA-treated group.

**Table 1.**
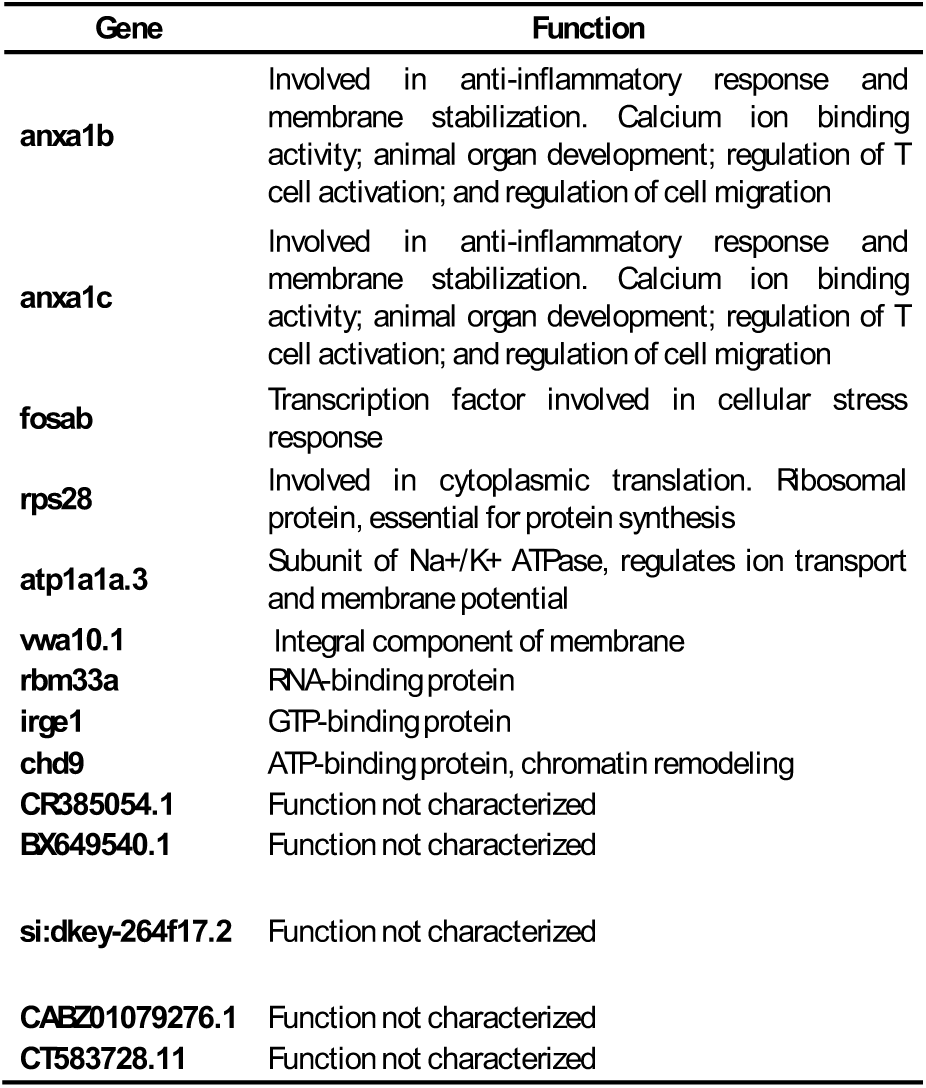
List of differentially expressed genes identified in zebrafish embryos treated with GBA 26 compared to water control, along with their known or predicted functions provided by NCBI Gene database (https://www.ncbi.nlm.nih.gov/gene/).

The annexin family of proteins can bind to cellular membranes in a calcium-dependent manner and is known to be involved in a variety of cellular processes, such as membrane trafficking, cell signalling, and particularly, the regulation of inflammation and response to cellular stress [47]. While anxa1b and anxa1c belong to the same family, they may have distinct or even opposing roles in certain cellular contexts, especially in response to environmental stress or chemical exposure [48].

The downregulation of anxa1b could indicate that this annexin isoform is suppressed in response to GBA and TPs, possibly due to reduced demand for its specific functions or because the cell is prioritizing other stress responses. This could be part of a broader adjustment to modulate the cell’s response to the biocide, potentially shifting to alternative pathways or isoforms, such as anxa1c. On the other hand, the upregulation of anxa1c suggests an active role in the cellular response to GBA. Anxa1c could be compensating for the downregulation of anxa1b, taking on more prominent roles in processes such as membrane repair or calcium signalling, both of which are crucial in stress responses [49]. Alternatively, anxa1c might have distinct functions specifically activated in response to GBAand TPs, such as facilitating stress-induced signaling pathways or regulating the inflammatory response.

Additional transcriptional changes included the downregulation of chd9 and fosab, linked to chromatin remodelling and stress-responsive transcription, respectively, and the upregulation of atp1a1a.3, encoding a Na⁺/K⁺-ATPase subunit involved in ion transport and membrane potential regulation [50]. The upregulation of rbm33a further suggests modulation of RNA processing and post-transcriptional regulation [51].

Collectively, these transcriptional changes reveal that GBA induces both upregulation and downregulation of key genes tied to stress response, ion transport, and transcriptional control. Rather than causing broad transcriptomic disruption, GBA appears to elicit targeted molecular adjustments, enhancing adaptive pathways to counter environmental stress. The observed upregulation of *anxa1c* and *atp1a1a.3*, alongside *anxa1b* downregulation, supports the idea that the response reflects cellular adaptation to GBA and TPs exposure rather than overt toxicity.

### 3.3 Toxicity to Raphidocelis subcapitata

Algae play a fundamental role in aquatic ecosystems, so disruptions to their growth can cascade through the food web. To assess the potential impact of GBA and DH345, an algal growth inhibition assay (OECD TG 201) was performed.

The results in Figures 8A-E illustrate the effects of GBA on *Raphidocelis subcapitata* growth over 72 hours.

Figure 8A illustrates a dose-dependent effect of GBA on algal growth over 72 hours. Biomass (cell/mL) increased substantially in the control and low concentrations (0.001, 0.01, and 0.05 mg/L), while higher concentrations (≥0.1 mg/L) significantly reduced biomass, despite all treatments showing exponential growth (>16-fold).

**Figure 8.**
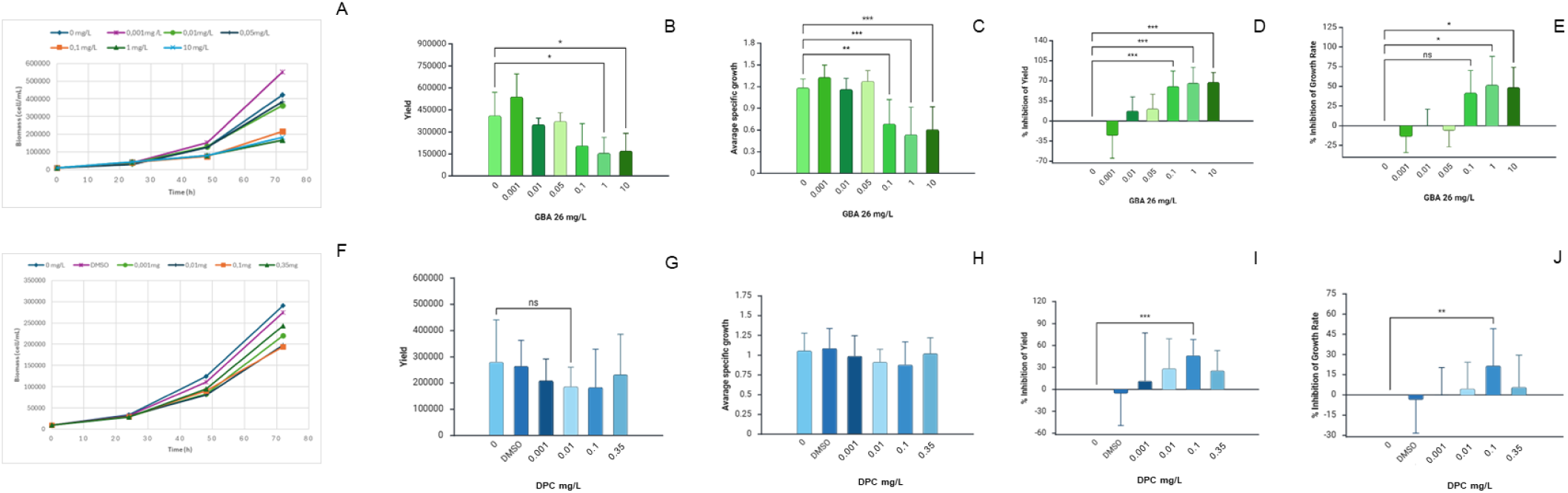
The effects of GBA 26 on algal growth (A-E) and DPC (F-J). (A, F) Biomass (cell/mL) measured at every 24h across a period of 72h. (B, G) Yield of cell density against different concentrations of GBA 26 (0 mg/L, 0.001 mg/L, 0.01 mg/L, 0.05 mg/L, 0.1 mg/L, 1 mg/L, and 10 mg/L) (C, H) Average specific growth rate against different concentrations of GBA 26 (D, I) Percentage of yield inhibition. (E, J) Percentage of growth rate inhibition. B, C, D: One-way-ANOVA (p&<0,05) with Dunnett post-hoc test. E: Krustal-Wallis Test (p&<0,05) with Bonferroni correction. Error bars represent standard deviations. * Indicates statistical significance (p &< 0.05); ** (p &< 0.01); *** (p &< 0.001).

This reduction is confirmed in Figure 8B, where yield (the difference between the initial and final biomass amounts) decreased markedly at 1 mg/L and above, with statistically significant differences from the control (p < 0.05). Lower concentrations caused no significant change.

Similarly, Figure 8C shows that the average specific growth rate followed the same trend, remaining high in the control and low dose treatments but dropping notably at ≥0.1 mg/L, indicating growth impairment likely due to cellular disruption.

An interesting observation is found in Figure 8D, which shows the percentage inhibition of yield. At ≥1 mg/L, inhibition is pronounced, but at the lowest concentration (0.001 mg/L), yield is slightly enhanced, resulting in negative inhibition values that are not significantly different from the control treatment. This observation suggests a possible hormetic response, where low doses of GB stimulate cellular processes or stress response mechanisms that temporarily enhance growth [52]. Finally, Figure 8E shows that the percentage inhibition of growth rate was significant at ≥1 mg/L.

Overall, these findings indicate that GBA only hinders algal growth at relatively high concentrations (≥0.1 mg/L), while low doses have minimal or no adverse effects, suggesting a concentration-dependent toxicity pattern.

The results in Figures 8F-J illustrate the effects of DH345 on *Raphidocelis subcapitata* growth over 72 hours.

Figure 8F shows that biomass increased exponentially across all treatments. In Figure 8G, a decreasing trend with concentration is present in yield, but without statistically significant differences. The average specific growth rate presented in Figure 8H also displayed no significant differences among treatments, indicating that overall growth dynamics remained stable within natural variability.

In contrast, the inhibition metrics displayed in Figures 8I and 8J revealed a dose-dependent response; both yield and growth rate inhibition increased significantly at 0.1 mg/L, but not at the highest concentration, and it is even possible to observe a slight stimulation at the lowest concentration, suggesting, again, a hormetic response.

Overall, both compounds show inhibition of algal growth at 0.1 mg/L, but GBA exhibits stronger effects at comparable doses. This comparison suggests that while DH345 poses a measurable risk to algal populations at higher concentrations, its impact is milder than that of GBA under equivalent conditions.

### 3.4 Toxicity to *Vibrio fischeri*

In this assay, *Vibrio fischeri* was exposed to three compounds (GBA, DH345, and tralopyril) and luminescence inhibition was measured.

GBA (10 mg/L) and DH345 (1 mg/L) caused no reduction in luminescence. Similarly, tralopyril, tested at 5 μg/L based on previous fish embryo test data, showed no luminescence inhibition. These results indicate that none of the compounds interfered with the bacterial respiratory process at the tested concentrations. Despite tralopyril’s high toxicity in zebrafish assays, these findings are consistent with previous studies [52], which also reported no effect on bacterial luminescence.

### 3.5 Transactivation assays: PPARy and PXR from *Mytilus galloprovincialis*

To investigate how NIAF interact with cellular receptors and influences gene expression, Nuclear Receptor transactivation assays were performed. NRs are transcription factors that initiate many physiological cascades regulating key physiological processes such as development, reproduction, and homeostasis. Most NRs are ligand-activated, making them both highly susceptible to disruption by foreign chemicals and, at the same time, crucial indicators of endocrine disruption.

In fact, the antifouling TBT serves as a textbook example of NR disruption. Common examples include compounds found in paints, detergents, that affect both the retinoid X receptor (RXR), causing imposex in gastropods [53], and the peroxisome proliferator-activated receptor gamma (PPAR), promoting adipogenesis [54].

To assess the potential endocrine-disrupting properties of GBA, preliminary studies evaluated its ability to modulate two nuclear receptors, in the model vertebrate *Danio rerio*: PPAR and Pregnane X Receptor (PXR) [23].

PPAR, predominantly expressed in adipose tissue and the intestine, plays a central role in adipocyte differentiation, lipid storage, and glucose homeostasis. It influences metabolic balance by regulating genes involved in lipid metabolism and insulin sensitivity through hormones such as adiponectin and leptin. Moreover, PPAR exhibits anti-inflammatory activity in metabolic tissues and may contribute to cardiovascular disease by modulating vascular function [55].

PXR, mainly expressed in the liver, is essential for xenobiotic metabolism (by cytochrome P450 regulation), detoxification, and lipid homeostasis. PXR is activated by a wide range of compounds due to its large and flexible ligand-binding domain [56].

Given that NR responses are not strictly conserved across taxa [57] and that these compounds exhibited antifouling activity mainly by decreasing *Mytilus galloprovincialis* settlement, the activity of PPAR and PXR receptors from *M. galloprovincialis* was further addressed.

Only GBA was tested since this NIAF appears to be the most promising, given the limited toxicity reported here. Previous studies indicated that GBA did not alter PPAR or PXR transcription in *D. rerio* [23]. To broaden phylogenetic coverage, additional assays were performed using *Mytilus galloprovincialis* NRs.

**Figure 9.**
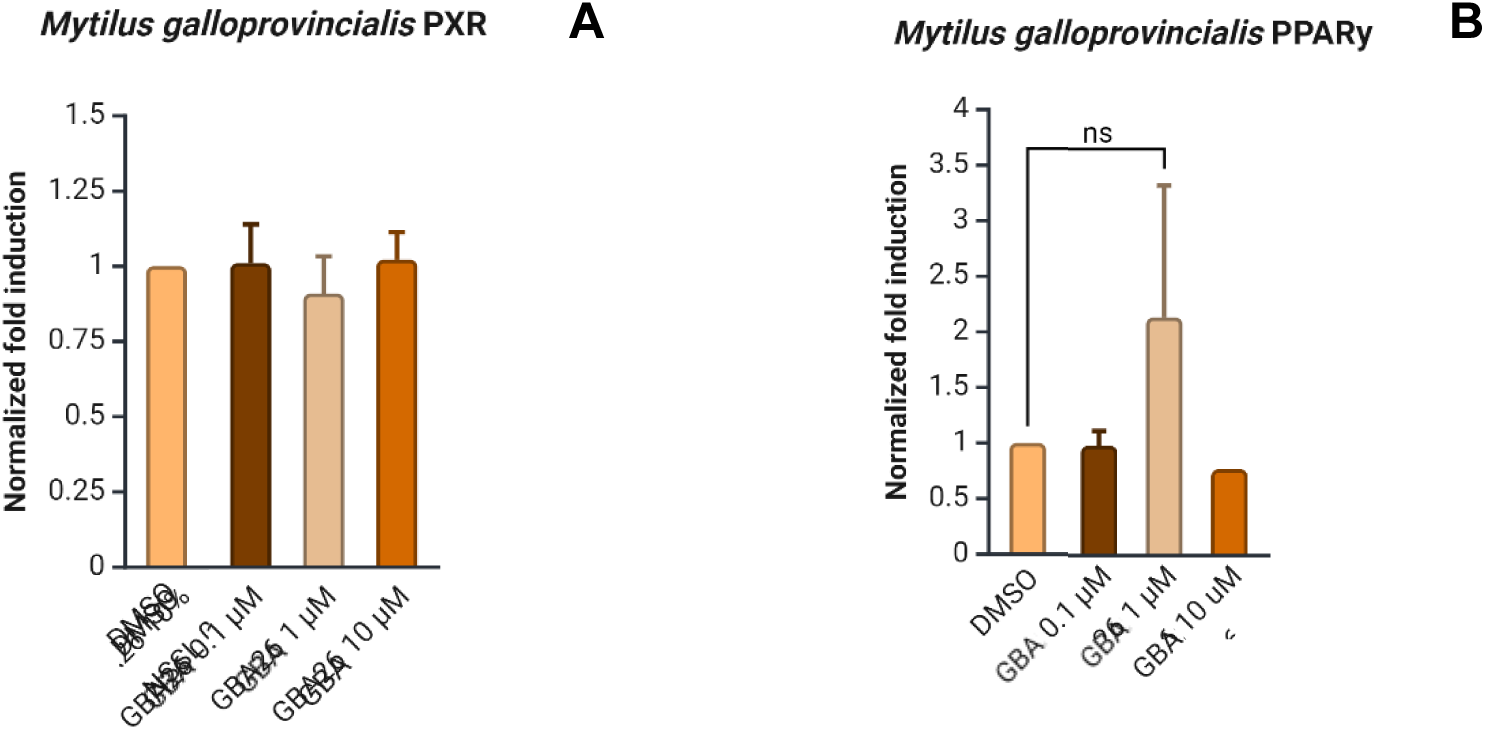
Effects of different treatments on nuclear receptor activation in *Mytilus galloprovincialis* measured by normalized fold induction of PXR (A) and PPARγ (B). (A) The bar graph shows the normalized fold induction of PXR in response to various treatments. No significant differences (ns) were observed between the treatments. **(B)** The bar graph illustrates the normalized fold induction of PPARγ in response to various treatments. No significant differences (ns) were detected between treatments. Treatments included different concentrations of GBA 26 (0.1, 1, and 10 μM) diluted in DMSO. Because these products were derived from a saltwater matrix, controls using equivalent saltwater dilutions (NSSL 0.1%, 1%, 10%) were also included. NSSL served as the control for each corresponding transformation product treatment (e.g. NSSL 0.1% for TP_GBA 26 0.1%), while DMSO served as the control for GBA 26 treatments. Krustal-Wallis Test (p&<0,05) with Bonferroni correction. Error bars represent standard deviations.

Figure 9A shows PPAR induction following exposure to DMSO, saltwater (NSSL) and GBA.

No statistically significant differences were detected; however, a slight, non-significant increase in fold induction was observed for GBA at 1 µM (≈2.6 mg/L), suggesting potential PPAR activation. Previous studies reported that exposure to *D. rerio* PPAR agonists and antagonists can reduce larval length and increase yolk sac area [58]; the latter effect was also observed in fish embryo tests (FET) with GBA at 1 mg/L. Nonetheless, given the higher variability observed at 1 µM, further analyses are needed to distinguish between biological response and replication variance.

Figure 9B presents the results for PXR induction under the same treatment conditions These findings indicate that GBA does not significantly affect PXR activity, a receptor central to the regulation of detoxification processes [59].

### 3.7 Determination of LOEC, NOEC and PNEC

Since the FET assay represents an acute test, an assessment factor (AF) of 1000 was applied to calculate the predicted no-effect concentrations (PNECs) for tralopyril. In contrast, for GBA and DH345, the algae growth inhibition test was identified as the most sensitive bioassay and was therefore used instead of the FET data. Although algal assays are typically considered acute, they can partially reflect chronic exposure since algae undergo multiple generations during the test period. For this reason, an AF of 100 is sometimes used [60]. However, to ensure a conservative and precautionary approach, an AF of 1000 was maintained, as indicated in Table 2.

**Table 2.** LOEC, NOEC, and PNEC values for three tested compounds (GBA, Tralopyril, and DPC). The table displays the Lowest Observed Effect Concentration (LOEC), No Observed Effect Concentration (NOEC), and Predicted No Effect Concentration (PNEC) for each compound. ¹Data from algae growth inhibition assay was used to determine values and an assessment factor of 1000 and 100 (denoted by an asterisk (*)) was applied. ² Data <u>from FET assay was used to determine values and an assessment factor of 1000 was applied.</u>

| Compound | LOEC µg/L | NOEC µg/L | PNEC µg/L |
| --- | --- | --- | --- |
| GBA<sup>1</sup> | 100 | 50 | 0.05/ 0.5* |
| DPC<sup>1</sup> | 100 | 10 | 0.01/0.1* |
| Tralopyril<sup>2</sup> | 1.03 | 0.515 | 0.000515 |

For GBA, the most sensitive endpoint was a NOEC of 50 μg/L in the algal assay, yielding a PNEC of 0.05 μg/L. For tralopyril, a NOEC could not be directly determined; thus, it was estimated by extrapolating from the LOEC (LOEC/2), yielding a PNEC of 0.515 ng/L. This aligns with literature findings reporting similar PNECs (e.g., 1.7 ng/L) from chronic Danio rerio studies with lower AF [61]. For DH345, a NOEC of 10 μg/L was established, leading to a PNEC of 0.01 μg/L.

## 4. CONCLUSION

Overall, GBA and DH345 showed no significant toxicity to zebrafish at 0.1 mg/L, while tralopyril was toxic at concentrations nearly 45 times lower, with a PNEC roughly 200 times lower. These findings suggest that, although GBA and DH345 exhibit some algal toxicity, they are considerably less toxic to vertebrate and bacterial models than tralopyril, reinforcing their potential as safer antifouling alternatives. Further chronic and multi-species studies are warranted to refine PNEC estimates and confirm their environmental safety profiles.

## Acknowledgments

This research was supported by FCT (Foundation for Science and Technology) within the scope of CIIMAR Base Funding UIDB/04423/2020 and UIDP/04423/2020 (DOI: 10.54499/ UIDB/04423/2020), UID/04423/2025 (DOI: 10.54499/UID/04423/2025), UID/PRR/04423/2025 (10.54499/UID/PRR/04423/2025) and LA/P/0101/2020 (10.54499/LA/P/0101/2020), and as a result of the project PTDC/CTA-AMB/0853/2021. Jessica Leite and Erica Lima acknowledge the scholarships supported by the NIAF project.

