## Supplementary material for "Eco-Friendly Antifouling Solutions: Hazard Assessment of Synthetic Derivatives of Natural Compounds": Suplementary material

[2]CIIMAR- Terminal de Cruzeiros de Leixões. AV. General Norton de Matos S/N 4450-208 Matosinhos

[3]Faculdade de Farmácia, Universidade do Porto, Rua de Jorge Viterbo Ferreira nº. 228, 4050-313, Porto, Portugal

Figure S1. A. Calibration curve of GBA in dechlorinated water. B. Chromatogram of the standards of the calibration curve of GBA hydrochloride in dechlorinated water.

Figure S2. Chromatogram and mass spectrum of fresh GBA26 standard in dechlorinated water.

Figure S3. Chromatogram and mass spectrum of 24h-GBA FET samples in dechlorinated water.

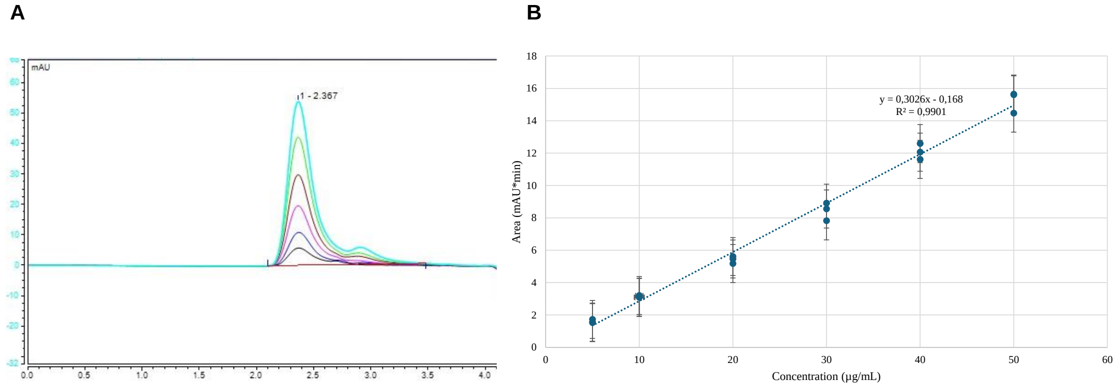


**Figure S1. A. Calibration curve of GBA26 in dechlorinated water. Standards: 5 µg/mL (black), 10 µg/mL (dark blue), 20 µg/mL (pink), 30 µg/mL (brown), 40 µg/mL (green) and 50 µg/mL (light blue). B. Chromatogram of the standards of the calibration curve of GBA26 in dechlorinated water.**

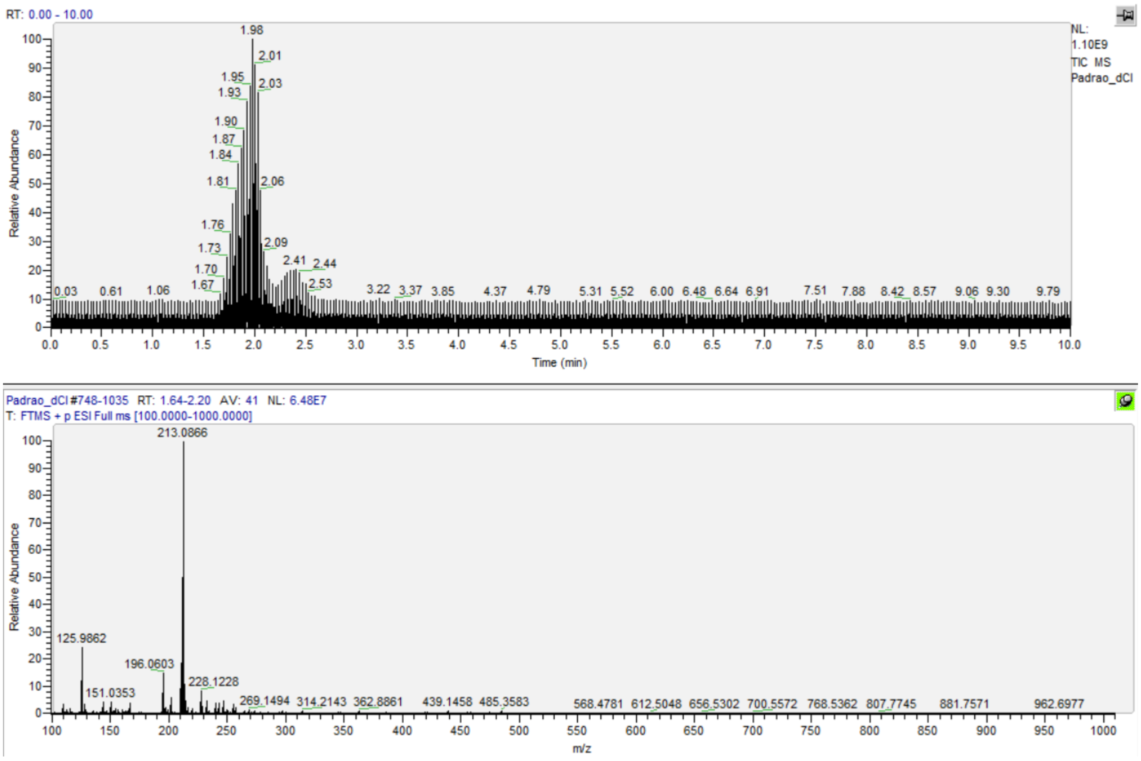


**Figure S2. Chromatogram and mass spectrum of fresh GBA26 standard in dechlorinated water.**

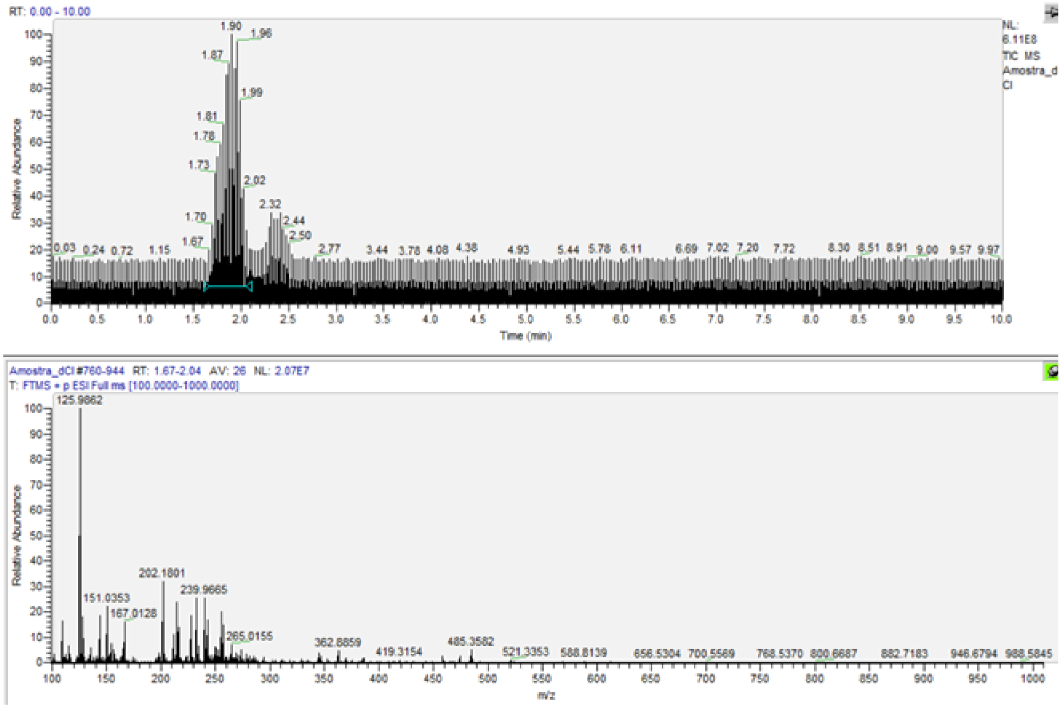


**Figure S3. Chromatogram and mass spectrum of 24h-GBA FET samples in dechlorinated water.**
